# GCA: An R package for genetic connectedness analysis using pedigree and genomic data

**DOI:** 10.1101/696419

**Authors:** Haipeng Yu, Gota Morota

**Author notes:** Corresponding author: Haipeng Yu, Department of Animal and Poultry Sciences, Virginia Polytechnic Institute and State University, 175 West Campus Drive, Blacksburg, Virginia 24061 USA.,; Gota Morota, Department of Animal and Poultry Sciences, Virginia Polytechnic Institute and State University, 175 West Campus Drive, Blacksburg, Virginia 24061 USA.

## Abstract

**Background:** Genetic connectedness is a critical component of genetic evaluation as it assesses the comparability of predicted genetic values across units. Genetic connectedness also plays an essential role in quantifying the linkage between reference and validation sets in whole-genome prediction. Despite its importance, there is no user-friendly software tool available to calculate connectedness statistics.

**Results:** We developed the GCA R package to perform genetic connectedness analysis for pedigree and genomic data. The software implements a large collection of various connectedness statistics as a function of prediction error variance or variance of unit effect estimates. The GCA R package is available at GitHub and the source code is provided as open source.

**Conclusions:** The GCA R package allows users to easily assess the connectedness of their data. It is also useful to determine the potential risk of comparing predicted genetic values of individuals across units or measure the connectedness level between training and testing sets in genomic prediction.

## Background

Genetic connectedness quantifies the extent to which estimated breeding values can be fairly compared across units or contemporary groups [1, 2]. Genetic evaluation is known to be more robust when the connectedness level is high enough due to sufficient sharing of genetic material across groups. In such scenarios, the best linear unbiased prediction minimizes the risk of uncertainty in ranking of individuals. On the other hand, limited or no sharing of genetic material leads to less reliable comparisons of genetic evaluation methods [3]. High-throughput genetic variants spanning the entire genome available for a wide range of agricultural species have now opened up an opportunity to assess connectedness using genomic data. A recent study showed that genomic relatedness strengthens the measures of connectedness across units compared with the use of pedigree relationships [4]. The concept of genetic connectedness was later extended to measure the connectedness level between reference and validation sets in whole-genome prediction [5]. This approach has also been used to optimize individuals constituting reference sets [6, 7] In general, it was observed that increased connectedness led to increased prediction accuracy of genetic values evaluated by cross-validation [8]. Comparability of total genetic values across units by accounting for additive as well as non-additive genetic effects has also been investigated [9].

Despite the importance of connectedness, there is no user-friendly software tool available that offers computation of a comprehensive list of connectedness statistics using pedigree and genomic data. Therefore, we developed a genetic connectedness analysis R package, GCA, which measures the connectedness between individuals across units using pedigree or genomic data. The objective of this article is to describe a large collection of connectedness statistics implemented in the GCA package, overview the software architecture, and present several examples using simulated data.

## Connectedness statistics

A list of connectedness statistics supported by the GCA R package is shown in Figure 1. These statistics can be classified into core functions derived from either prediction error variance (PEV) or variance of unit effect estimates (VE). PEV-derived metrics include prediction error variance of differences (PEVD), coefficient of determination (CD), and prediction error correlation (r). Further, each metric based on PEV can be summarized as the average PEV within and across units, at the unit level as the average PEV of all pairwise differences between individuals across units, or using a contrast vector. VE-derived metrics include variance of differences in unit effects (VED), coefficient of determination of VED (CDVED), and connectedness rating (CR). For each VE-derived metric, three correction factors accounting for the number of fixed effects can be applied. These include no correction (0), correcting for one fixed effect (1), and correcting for two or more fixed effects (2). Thus, a combination of core functions, metrics, summary functions, and correction factors uniquely characterizes connectedness statistics. Further, the overall connectedness statistic can be obtained by calculating the average of the pairwise connectedness statistics across units.

**Figure 1:**
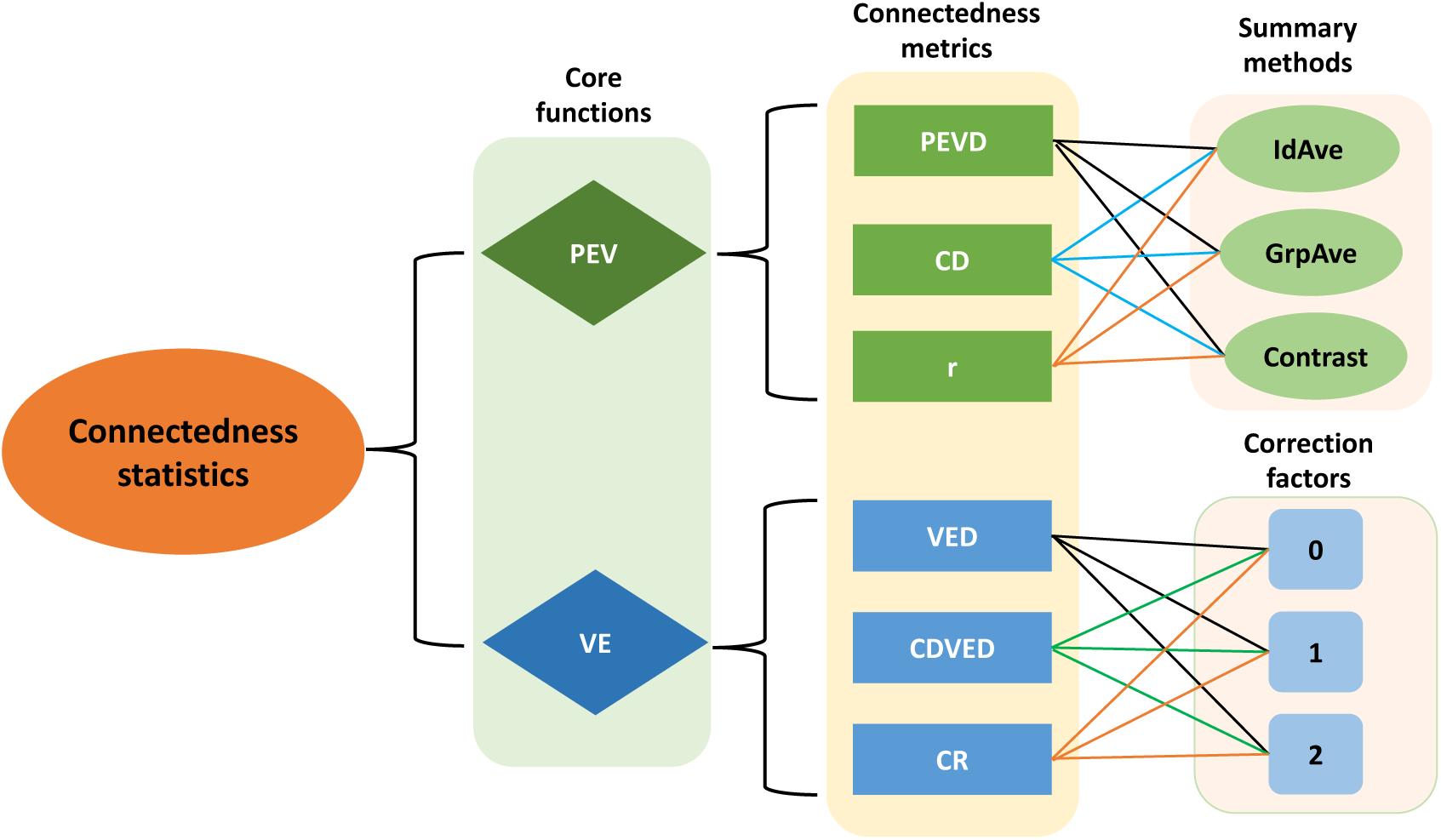
An overview of connectedness statistics implmented in the GCA R package. The statistics can be computed from either prediction error variance (PEV) or variance of unit effect estimates (VE). Connectedness metrics include prediction error variance of the difference (PEVD), coefficient of determination (CD), prediction error correlation (r), variance of differences in unit effects (VED), coefficient of determination of VE (CDVE), and connectedness rating (CR). IdAve, GrpAve, and Contrast correspond to individual average, group average, and contrast summary methods, respectively. 0, 1, and 2 are correction factors accounting for the fixed effects in the model.

### Core functions

#### Prediction error variance (PEV)

A PEV matrix is obtained from Henderson’s mixed model equations (MME) by assuming a standard linear mixed model **y** = **Xb** + **Zu** + ***ϵ***, where **y, b, u**, and ***ϵ*** refer to a vector of phenotypes, systematic effects, random additive genetic effects, and residuals, respectively [10]. The **X** and **Z** are incidence matrices associating systematic effects and genetic values to observations, respectively.

Then the PEV of **u** is derived as shown in Henderson [10].

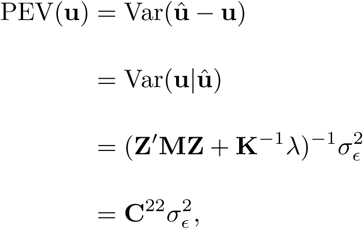

where **M** = **I** *-* **X**(**X**^*′*^**X**)^*-*^**X**^*′*^ is the absorption (projection) matrix for fixed effects, **K** is a relationship matrix, and 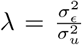 is the ratio of residual and additive genetic variance. **C**^22^ represents the lower right quadrant of the inverse of coefficient matrix. Note that PEV(**u**) can be viewed as the posterior variance of **u**.

#### Variance of unit effect estimates (VE)

An alternative option for the choice of core function is to use VE, which is based on the variance-covariance matrix of estimated unit or contemporary group effects. Kennedy and Trus [11] argued that mean PEV over unit (PEV_Mean_) defined as the average of PEV between individuals within the same unit can be approximated by 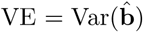, that is

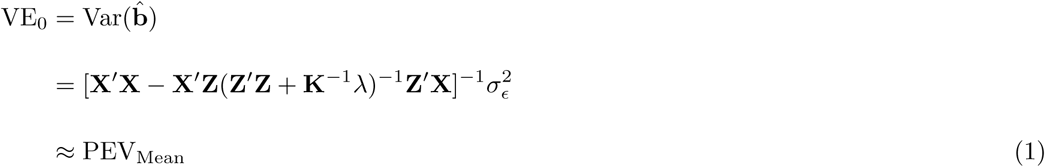

Holmes et al. [12] pointed out that the agreement between PEV_Mean_ and VE_0_ depends on a number of fixed effects other than the management group fitted in the model. They proposed exact ways to derive PEV_Mean_ as a function of VE and suggested addition of a few correction factors. When unit effect is the only fixed effect included in the model, the exact PEV_Mean_ can be obtained as given below.

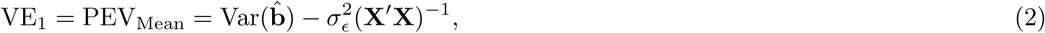

where **X**^*′*^**X**^*-*1^ is a diagonal matrix with *i*th diagonal element equal to 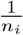, and *n*_*i*_ is the number of records in unit *i*. Thus, the term 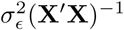 corrects the number of records within units. Accounting for additional fixed effects beyond unit effect when computing PEV_Mean_ is given by the following equation.

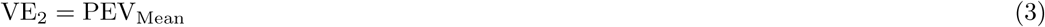

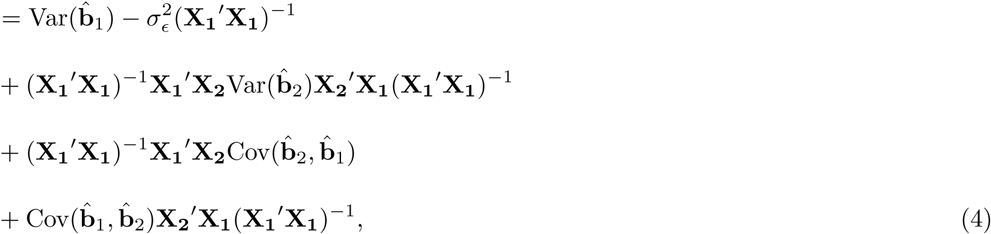

where **X**_**1**_ and **X**_**2**_ represent incidence matrices for units and other fixed effects, respectively, and 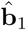 and 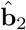 refer to the estimates of unit effects and other fixed effects, respectively [12]. This equation is suitable for cases in which there are two or more fixed effects fitted in the model.

### Connectedness metrics

Below we describe connectedness metrics implemented in the GCA package. We also summarized and organized their relationships with each other, which were never clearly articulated in the literature. These metrics are the function of PEV or VE described earlier (Figure 1).

#### Prediction error variance of difference (PEVD)

A PEVD metric measures the prediction error variance difference of breeding values between individuals from different units [11]. The PEVD between two individuals *i* and *j* is expressed as shown below.

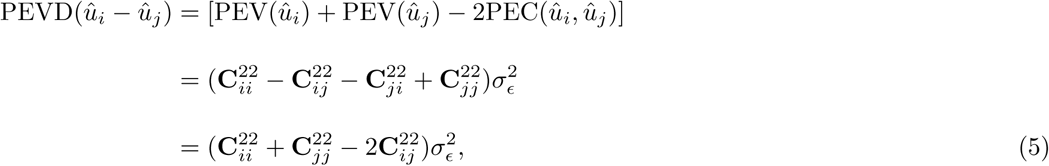

where PEC_*ij*_ is the off-diagonal element of the PEV matrix corresponding to the prediction error covariance between errors of genetic values.

##### Group average PEVD

The average PEVD derived from the average relationships between and within units as a choice of connectedness measure can be traced back to Kennedy and Trus [11]. This can be calculated by inserting the PEV_Mean_ of *i*^*′*^th and *j*^*′*^th units and mean prediction error covariance (PEC_Mean_) between *i*^*′*^th and *j*^*′*^th units into equation (5) as

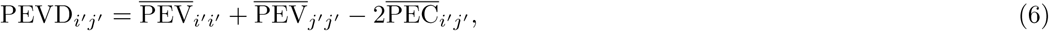

where 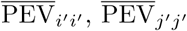, and 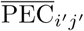 denote PEV_Mean_ in *i*^*′*^th and *j*^*′*^th units, and PEC_Mean_ between *i*^*′*^th and *j*^*′*^th units. We refer to this summary method as group average as illustrated in Figure 2A.

**Figure 2:**
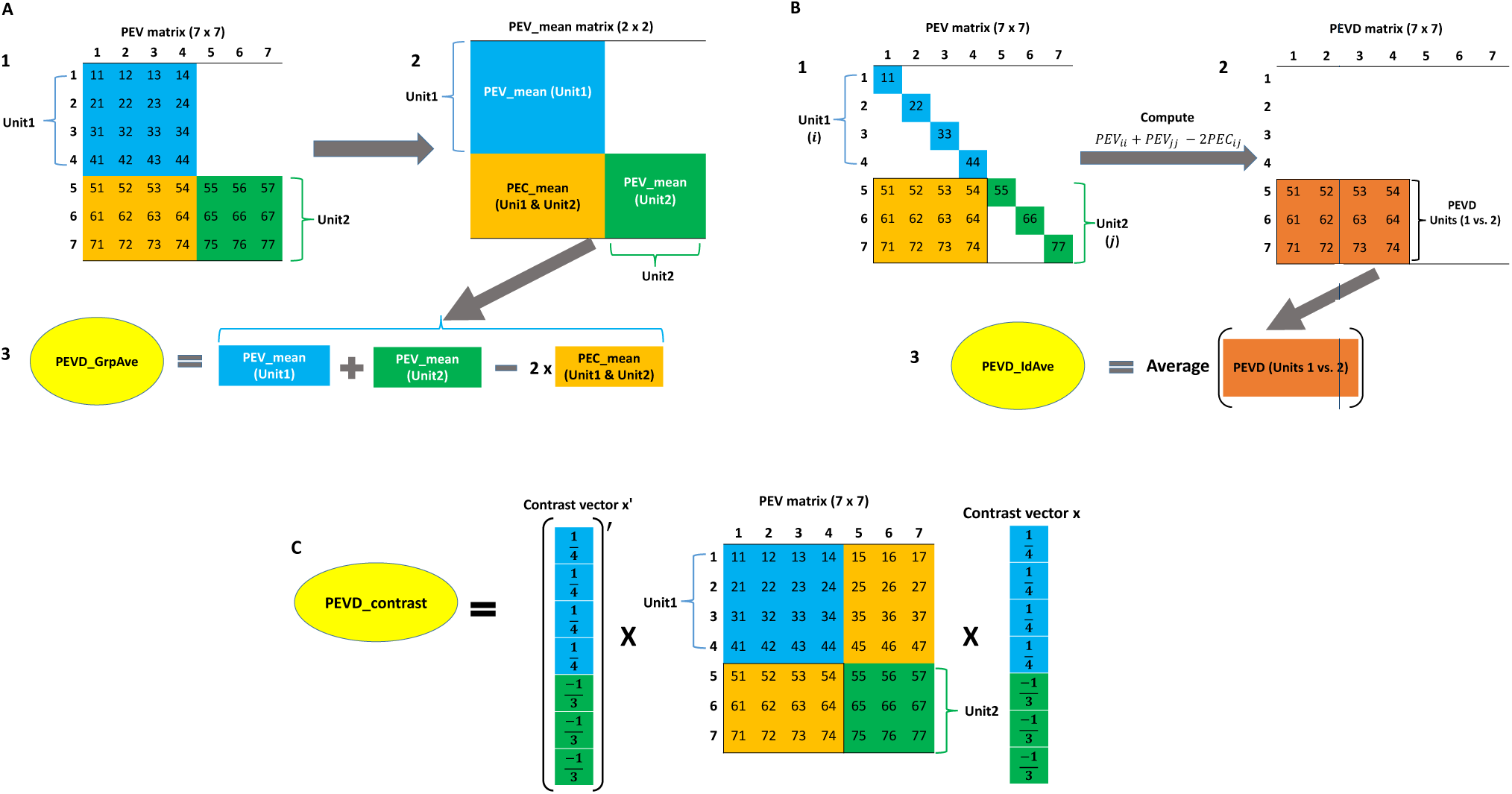
A flow diagram of three prediction error variance of the difference (PEVD) statistics. The group average PEVD (PEVD GrpAve) is shown in A. A1: Prediction error variance (PEV) matrix including variances and covariances of seven individuals. A2: Calculate the mean of prediction error variance / covariance within the unit (PEV mean) and mean of prediction error covariance across the unit (PEC mean). A3: Group average PEVD is calculated by applying the PEVD equation using PEV mean and PEC mean. The individual average PEVD (PEVD IdAve) is shown in B. B1: Prediction error variance (PEV) matrix including variances and covariances of seven individuals. Subscripts *i* and *j* refer to the *i*th and *j*th individuals in units 1 and 2, respectively. B2: Pairwise PEVD between individuals across two units. B3: Individual average PEVD is calculated by taking the average of all pairwise PEVD. The PEVD of contrast (PEVD Contrast) is shown in C. PEVD Contrast is calculated as the product of the transpose of the contrast vector (**x**), the PEV matrix, and the contrast vector.

##### Individual average PEVD

Alternatively, we can first compute PEVD at the individual level using equation (5) and then aggregate and summarize at the unit level to obtain the average of PEVD between individuals across two units [13]

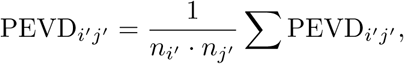

where *n*_*i*′_ and *n*_*j*′_ are the total number of records in units *i*^*′*^ and *j*^*′*^, respectively and ΣPEVD_*i*′*j′*_ is the sum of all pairwise differences between the two units. We refer to this summary method as individual average. A flow diagram illustrating the computational procedure is shown in Figure 2B.

##### Contrast PEVD

The PEVD of contrast between a pair of units can be used to summarize PEVD [14].

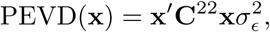

where **x** is a contrast vector involving 1*/n*_*i*′_, 1*/n*_*j*′_ and 0 corresponding to individuals belonging to *i*^*′*^th, *j*^*′*^th, and the remaining units. The sum of elements in **x** equals to zero. A flow diagram showing a computational procedure is shown in Figure 2C.

#### Coefficient of determination (CD)

A CD metric measures the precision of genetic values and can be interpreted as the square of the correlation between the predicted and the true difference in the genetic values or the ratio of posterior and prior variances of genetic values **u** [15]. A notable difference between CD and PEVD is that CD penalizes connectedness measurements when across units include individuals that are genetically too similar [4, 8]. A pairwise CD between individuals *i* and *j* is given by the following equation.

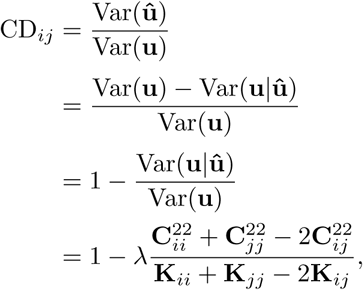

where **K**_*ii*_ and **K**_*jj*_ are *i*th and *j*th diagonal elements of **K**, and **K**_*ij*_ is the relationship between *i*th and *j*th individuals [14].

##### Group average CD

Similar to the group average PEVD statistic, PEV_Mean_ and PEC_Mean_ can be used to summarize CD at the unit level.

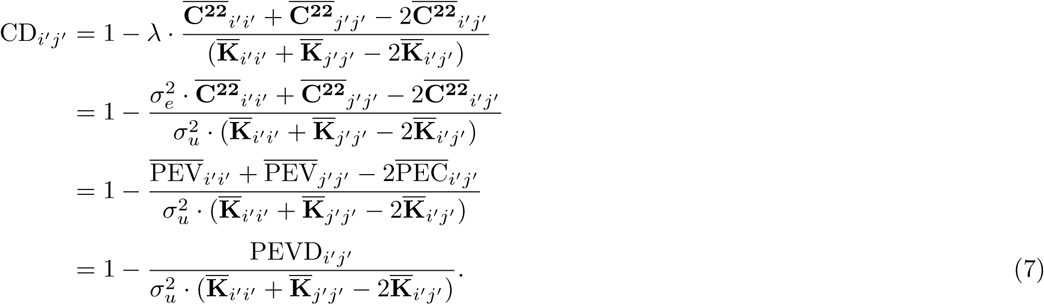

Here, 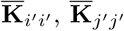 and 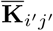 refer to the means of relationship coefficients in units *i*^*′*^ and *j*^*′*^, and the mean relationship coefficient between two units *i*^*′*^ and *j*^*′*^, respectively. Graphical derivation of group average CD is illustrated in Figure 3A. This summary method has not been used in the literature, but shares the same spirit with the group average PEVD.

**Figure 3:**
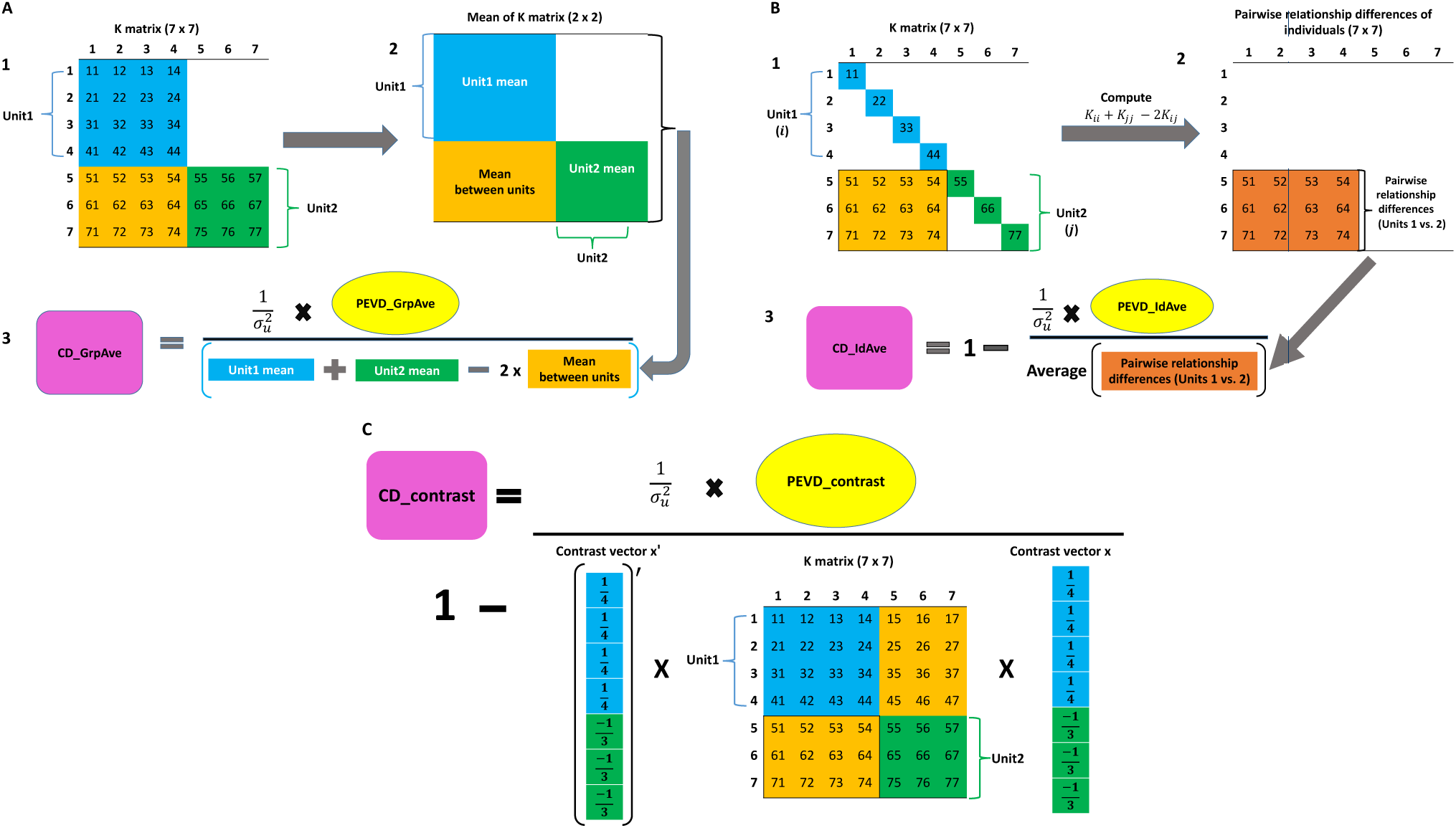
A flow diagram of three coefficient of determination (CD) statistics. The group average CD (CD GrpAve) is shown in A. A1: A relationship matrix of seven individuals. A2: Calculate the mean relationships within and between units. A3: Group average CD is calculated by scaling group average PEVD (PEVD GrpAve) by the quantity obtained from the PEVD equation using the within and between unit means. The individual average CD (CD IdAve) is shown in B. B1: A relationship matrix of seven individuals. B2: Calculate pairwise relationship differences of individuals between the units. Subscripts *i* and *j* refer to the *i*th and *j*th individuals in units 1 and 2, respectively. B3: Individual average CD is calculated by scaling indvidual average PEVD (PEVD IdAve) with the average of pairwise relationship differences of individuals. The CD of contrast (CD Contrast) is shown in C. CD Contrast is calculated by scaling the prediction error variance of the differences (PEVD) of contrast with the product of the transpose of the contrast vector (**x**), the relationship matrix (**K**), and the contrast vector.

##### Individual average CD

Individual average CD is derived from the average of CD between individuals across two units [13].

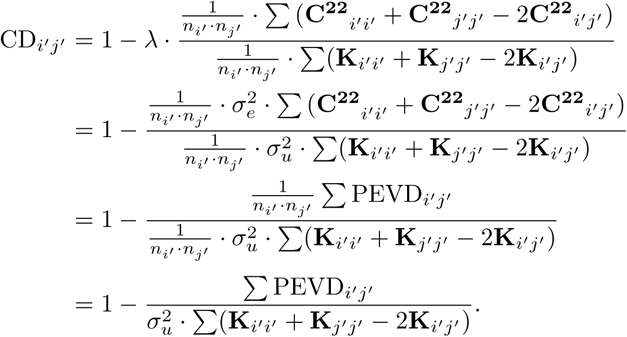

A flow diagram of individual average CD is shown in Figure 3B.

##### Contrast CD

A contrast of CD between any pair of units is given by [14]

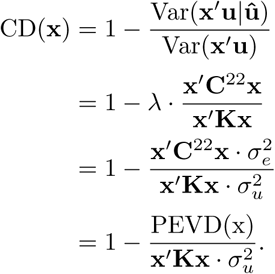

A flow diagram showing the computational procedure is shown in Figure 3C.

#### Prediction error correlation (r)

Prediction error correlation, known as pairwise r statistic, between individuals *i* and *j* is calculated from the elements of the PEV matrix [16].

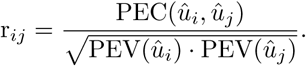

##### Group average r

This is known as flock connectedness in the literature, which calculates the ratio of PEV_Mean_ and PEC_Mean_ [16]. This group average connectedness for r between two units *i*^*′*^ and *j*^*′*^ is given by

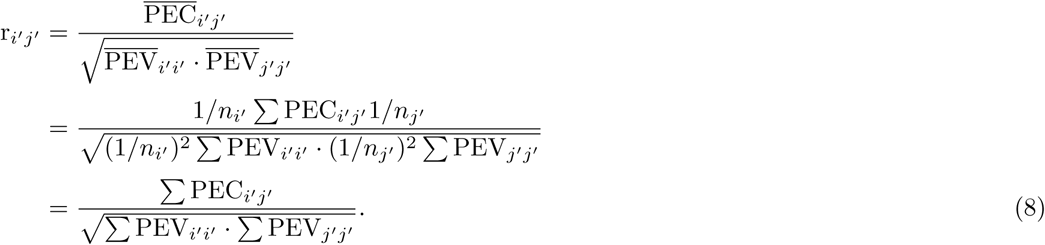

A graphical derivation is presented in Figure 4A.

**Figure 4:**
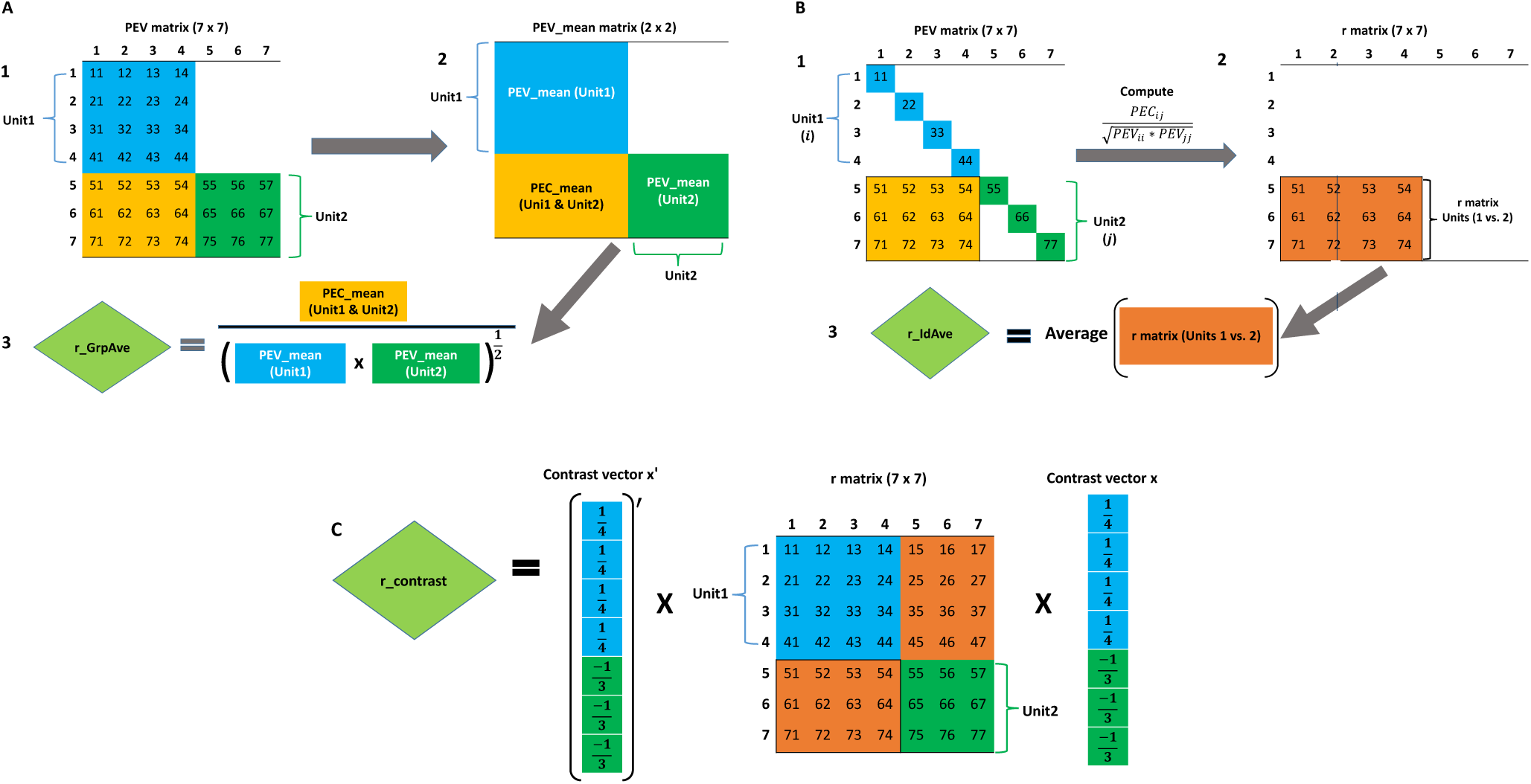
A flow diagram of three prediction error correlation (r) statistics. The group average r (r GrpAve) is shown in A. A1: Prediction error variance (PEV) matrix of seven individuals. A2: Calculate the mean of prediction error variance / covariance within the unit (PEV mean) and mean of prediction error covariance across the unit (PEC mean). A3: Group average r is a correlation calculated from PEV mean and PEC mean. The calculation of individual average r (r IdAve) involving seven individuals is displayed in B. B1: Prediction error variance (PEV) matrix of seven individuals. B2: Calculate pairwise correlation coefficients of individuals between units using PEV and prediction error covariance (PEC). Subscripts *i* and *j* refer to the *i*th and *j*th individuals in units 1 and 2, respectively. B3: Individual average r is calculated as the average of pairwise prediction error correlation coefficients of individuals across units. The r of contrast (r Contrast) is shown in C. r Contrast is calculated from the product of the transpose of the contrast vector (**x**), r matrix, and the contrast vector.

##### Individual average r

The summary method based on individual average calculates pairwise r for all pairs of individuals followed by averaging all r measures across units.

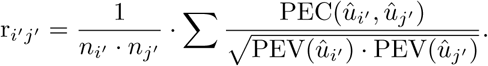

This summary method was first used in Yu et al. [4] and calculation steps are shown in Figure 4B.

##### Contrast r

A contrast of r is defined as below.

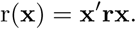

This summary method has not been used in the literature, but shares the same concept with the contrasts PEVD and CD. A flow diagram illustrating a computational procedure is shown in Figure 4C.

### Variance of differences in unit effects (VED)

A metric VED, which is a function of VE can be used to measure connectedness. All PEV-based metrics follow a two-step procedure in the sense that they first compute the PEV matrix at the individual level and then apply one of the summary methods to derive connectedness at the unit level or vice versa. In contrast, VE-based metrics follow a single-step procedure such that we can obtain connectedness between units directly. Moreover, since the number of fixed effects is oftentimes smaller than the number of individuals in the model, the computational requirements for VED are expected to be lower [12]. Note that all VE-derived approaches can be classified based on the number of fixed effects to be corrected. Using the group average summary method, three VEDc statistics estimate PEVD alike connectedness between two units *i*^*′*^ and *j*^*′*^ by replacing PEV_Mean_ in equation (6) from VEc [11, 12].

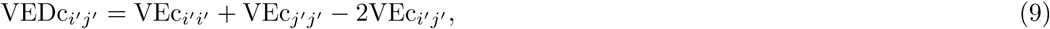

Here, c denotes no correction (0), correction for one fixed effect (1), and correction for two or more fixed effects (2) [12].

#### Coefficient of determination of VED

Similarly, the correction function based on VEc can be employed to define a group average CD alike statistic. We named this as CDVEDc. A pairwise CDVEDc between two units *i*^*′*^ and *j*^*′*^ is given by

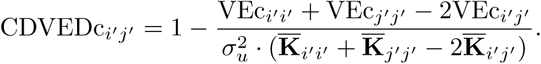

Here, c includes 0, 1, and 2 by referring to the number of corrections for fixed effects.

#### Connectedness rating (CR)

A CR statistic first proposed by Mathur et al. [17] is similar to equation (8). However, it uses variances and covariances of estimated unit effects instead of PEV_Mean_ and PEC_Mean_. Holmes et al. [12] extended CR by replacing VE with VEc to calculate CR and this is referred as CRc below. A pairwise CRc between two units *i*^*′*^ and *j*^*′*^ is outlined as

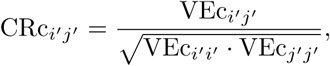

where c equals to the number of corrections for fixed effects: 0, 1, and 2. When c is set to 0, this is equivalent to CR of Mathur et al. [17].

### Software Description

#### Overview of software architecture

The GCA R package is implemented entirely in R, which is an open source programming language and environment for performing statistical computing [18]. The package is hosted on a GitHub page accompanied by a detailed vignette document. Computational speed was improved by integrating C++ code into R code using the Rcpp package [19]. The initial versions of the algorithms and the R code were used in previous studies [4, 8, 9] and were enhanced further for efficiency, usability, and documentation in the current version to facilitate connectedness analysis. The GCA R package provides a comprehensive and effective tool for genetic connectedness analysis and whole-genome prediction, which further contributes to the genetic evaluation and prediction.

### Installing the GCA package

The current version of the GCA R package is available at GitHub (https://github.com/QGresources/GCA). The package can be installed using the devtools R package [20] and loaded into the R environment following the steps shown at GitHub.

### Simulated data

A simulated cattle data set using QMSim software [21] is included in the GCA package as an example data set. A total of 2,500 cattle spanning five generations were simulated with pedigree and genomic information available for all individuals. We simulated 10,000 evenly distributed biallelic single nucleotide polymorphisms and 2,000 randomly distributed quantitative trait loci (QTL) across 29 pairs of autosomes with 100 cM per chromosome. A single phenotype with a heritability of 0.6 and a fixed covariate of sex were simulated. This was followed by simulating units using the k-medoid algorithm [22] coupled with the dissimilarity matrix derived from a numerator relationship matrix as shown in previous studies [4, 8, 9]. The data set is stored as an R object in the package. The genotype object is a 2, 500 *×* 10, 000 marker matrix. The phenotype object is a 2, 500 *×* 6 matrix, including the columns of progeny, sire, dam, sex, unit, and phenotype.

### Application of the GCA package

A detailed usage of the GCA R package can be found in the vignette document (https://qgresources.github.io/GCA_Vignette/GCA.html). Examples include 1) pairwise and overall connectedness measures across units; 2) relationship between PEV- and VE-based connectedness metrics; and 3) relationship between connectedness metrics and genomic prediction accuracies.

## Conclusions

The GCA R package provides users with a comprehensive tool for analysis of genetic connectedness using pedigree and genomic data. The users can easily assess the connectedness of their data and be mindful of the uncertainty associated with comparing genetic values of individuals involving different management units or contemporary groups. Moreover, the GCA package can be used to measure the level of connectedness between training and testing sets in the whole-genome prediction paradigm. This parameter can be used as a criterion for optimizing the training data set. This paper also summarized the relationship among various connectedness metrics, which was not clearly articulated in the past literature. In summary, we contend that the availability of the GCA package to calculate connectedness allows breeders and geneticists to make better decisions on comparing individuals in genetic evaluations and inferring linkage between any pair of individual groups in genomic prediction.

## Availability and implementation

The GCA R source code is provided as free and open source. The webpage https://github.com/QGresources/GCA was created as a nexus of all genetic connectedness related functions and examples available in the GCA R package. The vignette is available at https://qgresources.github.io/GCA_Vignette/GCA.html.

## Authors’ contributions

HY and GM developed the software tool and wrote the manuscript. GM supervised and directed the study. All authors read and approved the manuscript.

### Ethics approval and consent to participate

Not applicable.

### Consent for publication

Not applicable.

